# Total cell N-glycosylation is altered during neuronal differentiation of iPSC to NSC and is disturbed by trisomy 21

**DOI:** 10.1101/2023.06.28.546867

**Authors:** Ana Cindrić, Frano Vučković, Aoife Murray, Thomas Klarić, Ivan Alić, Dražen Juraj Petrović, Jasminka Krištić, Dean Nižetić, Gordan Lauc

## Abstract

Down syndrome (DS) is a genetic condition caused by trisomy 21 (T21) and characterized by a range of neurological symptoms including intellectual disability, early onset of neurodegeneration and dementia, some of which can be attributed to accelerated aging. N-glycosylation is a protein modification that plays a critical role in numerous biological processes and its dysregulation is associated with a wide range of diseases, in some even before the onset of symptoms. N-glycosylation of total plasma proteins, as well as specific plasma proteins, such as immunoglobulin G, has been shown to change in DS, displaying an accelerated aging phenotype consistent with the various symptoms of premature aging that occur in DS. However, little is known about how T21 affects the N-glycosylation of other cellular proteins. To better understand how T21 affects N-glycosylation during neural differentiation, we characterized and compared the total released N-glycans of induced pluripotent stem cells (iPSCs) and their neural stem cell (NSC) derivatives. We analyzed six different isogenic clones all derived from a single individual with mosaic DS and thus all sharing the same genetic background; however, three had a normal disomic karyotype (D21), while the other three had an additional copy of chromosome 21 (T21). We characterized the total cell N-glycosylation profiles using ultra high performance liquid chromatography (UHPLC) and subsequent tandem mass spectrometry analysis to determine proposed glycan structures. Our results revealed both qualitative and quantitative differences in the composition of N-glycomes between iPSCs and NSCs, with NSCs showing a higher amount of complex N-glycans and a lower amount of mannosidic N-glycans when compared to iPSCs. Moreover, we found differences in N-glycosylation patterns between D21 and T21 cells. Notably, T21 cells exhibited a significant increase in the amount of pseudohybrid N-glycans. Trisomy 21 also caused a significant decrease in the abundance of a hybrid monoantennary fucosylated glycan (H6N3F1). Our findings define the released N-glycan profile of total cells for both D21 and T21 iPSCs and NSCs and suggest that the presence of a third copy of chromosome 21 impacts N-glycosylation patterns already in the stem cell state.

## Introduction

N-glycosylation is a co- and posttranslational protein modification crucial for nearly every known physiological process (Varki & Kornfeld, 2022). N-glycans all share a conserved core comprised of two N-acetylglucosamine (GlcNAc) residues and three mannose (Man) residues. Depending on what type of monosaccharides are added to the core, they can be divided into three main subgroups ranging from the least to most processed: oligomannose, in which the core is elongated only by Man residues; hybrid, in which the core is modified by a combination of Man residues and GlcNAc-initiated antennae; and complex N-glycans, in the core is extended by two or more GlcNAc-initiated antennae (Varki et al., 2022). On a cellular level, glycosylation is important for processes such as cell migration, adhesion, signalling, differentiation, and immune recognition (Gu & Taniguchi, 2008; Ohtsubo & Marth, 2006; van Kooyk & Rabinovich, 2008). Aberrations in glycosylation have been linked to a range of diseases, including cancer, autoimmune disorders, and genetic disorders (Lauc & Trbojević-Akmačić, 2021; Varki et al., 2022). In the nervous system, N-glycosylation is dynamically regulated (Klarić & Lauc, 2022) and has been proven to be crucial for proper development and function (Conroy et al., 2021; Scott & Panin, 2014).

Down syndrome (DS) is a condition caused by full or partial trisomy 21 (T21) (Antonarakis et al., 2020; Lejeune, Jerome; Gautier, M; Turpin, 1959), and it presents with various comorbidities, many of which can be attributed to accelerated ageing (Head et al., 2012; Horvath et al., 2015; Meharena et al., 2022; Raji & Rao, 1998) and tend to affect the nervous system, potentially contributing to intellectual disability and dementia (Wiseman et al., 2015). The DS critical region (DSCR) is a segment on chromosome 21 implied to contain genes responsible for numerous features of DS (Pelleri et al., 2016; Shapiro, 1999). Among the ∼30 genes in the DSCR, only one gene is directly involved in glycosylation*–B3GALT5*, a gene encoding a beta-1,3-galactosyl transferase whose preferential substrates are glycolipids and O-glycans. Still, it has been shown that N-glycosylation of total plasma proteins is altered in DS (Borelli et al., 2015), wherein the shift in N-glycosylation is consistent with an accelerated ageing phenotype known to occur in DS (Borelli et al., 2015; Cindrić et al., 2021). In the study of IgG N-glycans, it was indicated that only a 31-gene segmental duplication that contains the entire DSCR is both sufficient and responsible for the IgG N-glycome shift toward accelerated ageing (Cindrić et al., 2021).

We have previously developed an isogenic induced pluripotent stem cell (iPSC) model of DS in which both D21 and T21 clones were derived from a single adult with mosaic DS to study the isolated effect of T21 excluding all other genetic variables. This model has several advantages over other cellular DS models: it is a fully human model that is non-integrationally reprogrammed, fully isogenic, and does not suffer from artificial genomic rearrangements. Neurons derived from the T21 iPSCs show several cellular pathologies linked with DS, reproducing DS-characteristic abnormalities in cell differentiation, aging, and neurodegeneration (Murray et al., 2015).

To the best of our knowledge, no previous studies have examined the effect of T21 on N-glycan profiles of human iPSCs or neural stem cells (NSCs). N-glycosylation changes occurring during differentiation from human iPSCs to NSCs have been studied on a glycopeptide level using mass spectrometry (MS), and only on one 46XX (normal karyotype) cell line (Kimura et al., 2021). In this study we characterize and compare total released N-glycans from iPSCs and their NSC derivatives through analysis of six different isogenic clones derived from the same person with mosaic DS. Three of the clones have a 46XX karyotype while the other three have an extra copy of chromosome 21. We offer a comparison of N-glycosylation during cellular differentiation from iPSCs to NSCs, as well as a comparison of N-glycosylation between isogenic cells that only differ by an additional copy of chromosome 21 and uncover some underlying differences using ultra high performance liquid chromatography (UHPLC) and subsequent tandem MS analysis.

## Results

### Cell validation/characterization

iPSCs were validated by immunostaining to confirm the expression of a panel of pluripotency markers (OCT4, SSEA4, TRA1-60 and TRA1-81) (**Supplementary Figure 1A**). NSCs were also validated by immunostaining to confirm the expression of Nestin, SOX2 and PAX6 (**Supplementary Figure 1B**). In addition, disomy/trisomy 21 of the cells was confirmed by FISH.

### N-glycosylation analysis

We analyzed the total cell N-glycosylation profile of isogenic iPSCs, all derived from the same individual with mosaic DS (n = 6) and their NSC derivatives (n = 6). Three of the six clonal cell lines were disomic for chromosome 21 (D21), while the remaining three had trisomy 21 (T21). The relative abundance of individual N-glycan peaks (GPs) and, thus, the overall N-glycosylation profiles of the cells were determined by UHPLC while qualitative structural analysis of representative pooled samples from each experimental group was performed by tandem MS to assign proposed N-glycan structures to each chromatographic peak (**Figure 1, Table 1, Supplementary Table 1**). A comparison of N-glycomes was done both on a qualitative and quantitative level.

**Figure 1.**
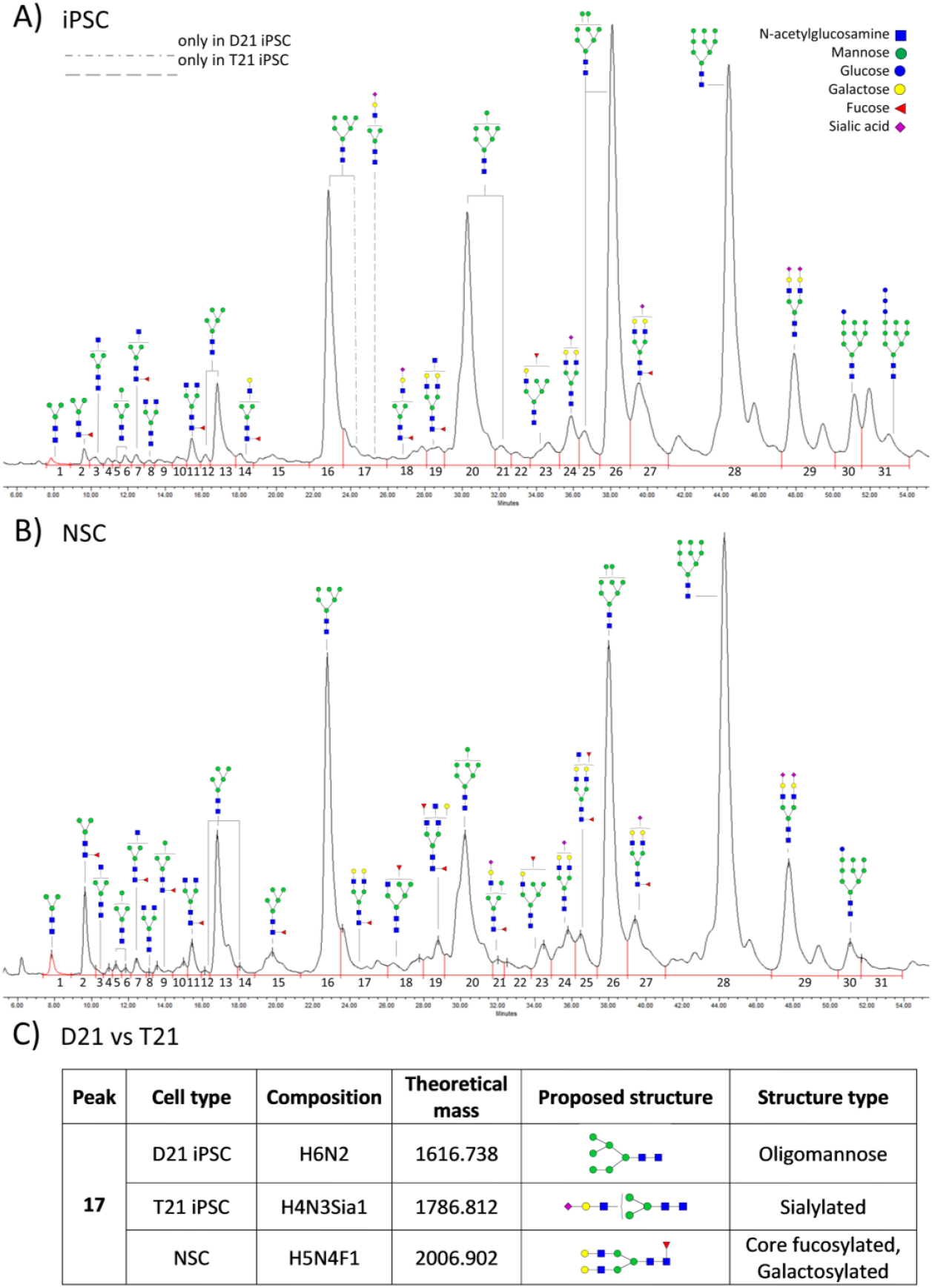
Released N-glycans of both analyzed cell types. A) Annotated chromatogram of induced pluripotent stem cell (iPSC), and B) neural stem cell (NSC). The most abundant structure from each glycan peak is shown where possible., C) A closer look at GP17, a chromatographic peak whose abundance differs between iPSCs with disomy 21 (D21) and trisomy 21 (T21).

**Table 1.**
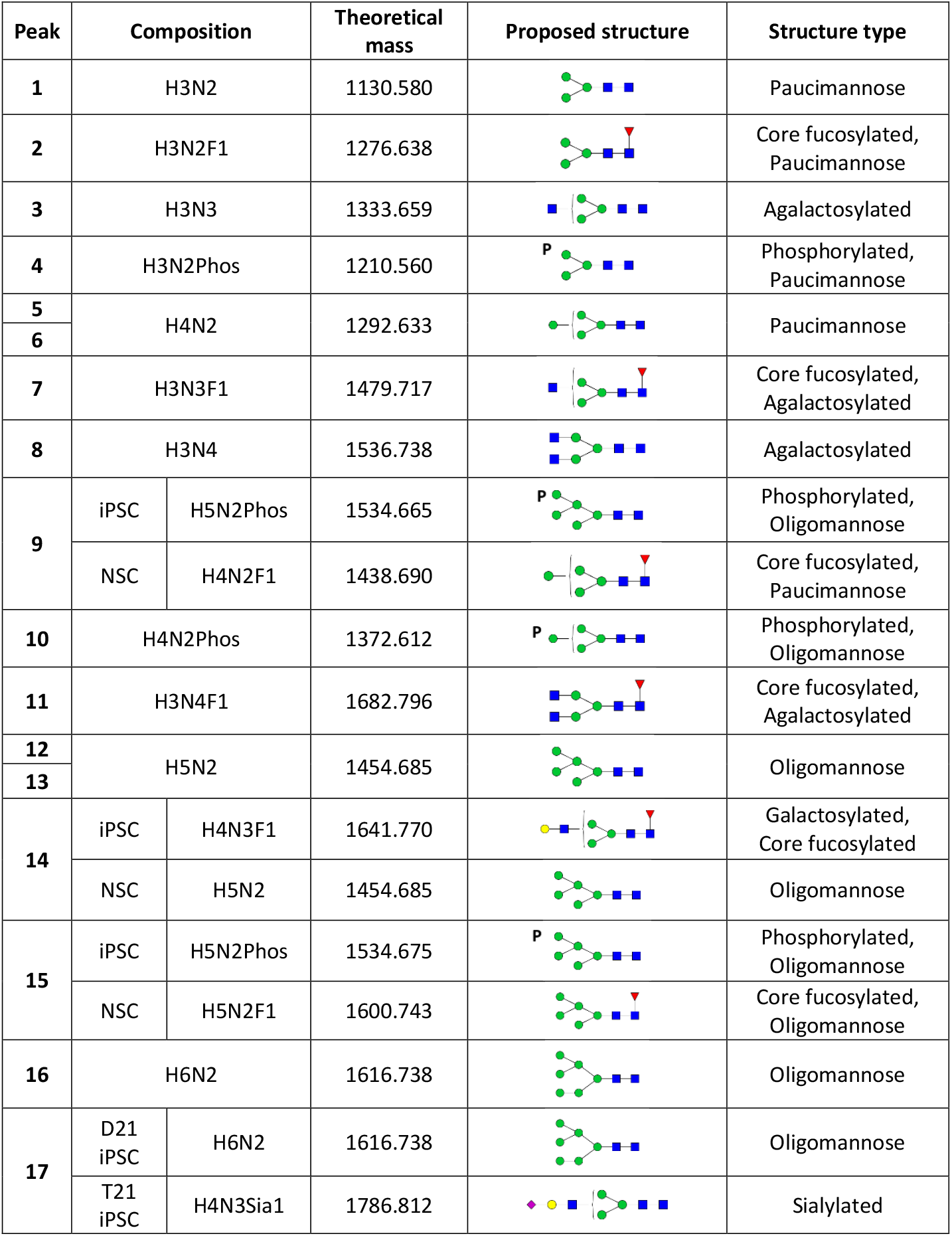
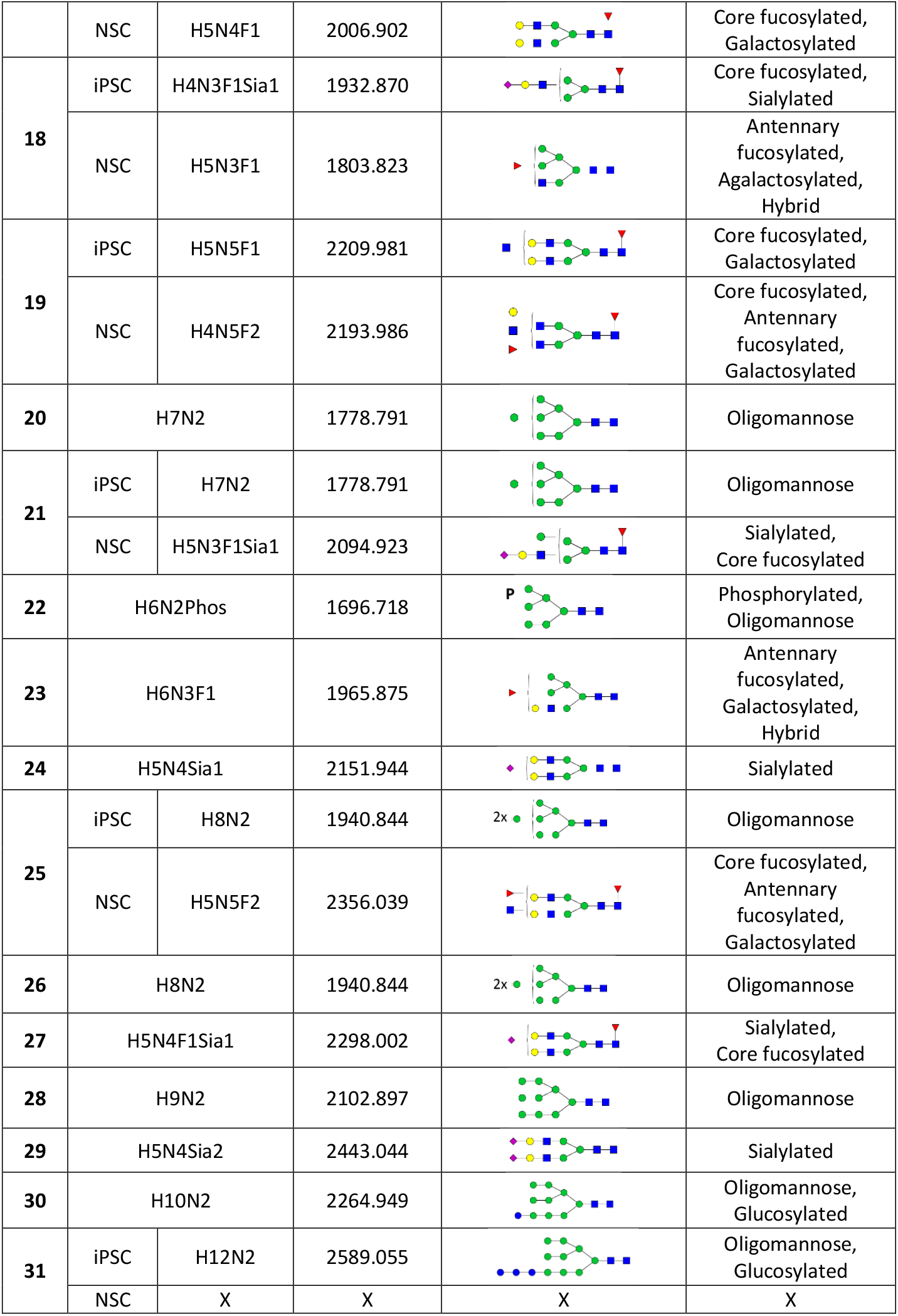
The most abundant glycan masses and corresponding composition and proposed structure per peak. The structure types have been divided into pauci- and oligomannose type glycans, phosphorylated glycans, and glycans containing fucose (core and antennary), galactose (a/galactosylated), and sialic acid residue/s. H – hexose, N – N-acetylglucosamine, F – fucose, Phos – phosphate, Sia – sialic acid, iPSC – induced pluripotent stem cells, NSC – neural stem cells, D21 – disomic cells, T21 – cells with three copies of chromosome 21, X – no structure was detected.

| Peak | Composition |  | Theoretical mass | Proposed structure | Structure type |
| --- | --- | --- | --- | --- | --- |
| 1 | H3N2 |  | 1130.580 |  | Paucimannose |
| 2 | H3N2F1 |  | 1276.638 |  | Core fucosylated, Paucimannose |
| 3 | H3N3 |  | 1333.659 |  | Agalactosylated |
| 4 | H3N2Phos |  | 1210.560 |  | Phosphorylated, Paucimannose |
| 5 | H4N2 |  | 1292.633 |  | Paucimannose |
| 6 |  |  |  |  |  |
| 7 | H3N3F1 |  | 1479.717 |  | Core fucosylated, Agalactosylated |
| 8 | H3N4 |  | 1536.738 |  | Agalactosylated |
| 9 | iPSC | H5N2Phos | 1534.665 |  | Phosphorylated, Oligomannose |
|  | NSC | H4N2F1 | 1438.690 |  | Core fucosylated, Paucimannose |
| 10 | H4N2Phos |  | 1372.612 |  | Phosphorylated, Oligomannose |
| 11 | H3N4F1 |  | 1682.796 |  | Core fucosylated, Agalactosylated |
| 12 | H5N2 |  | 1454.685 |  | Oligomannose |
| 13 |  |  |  |  |  |
| 14 | iPSC | H4N3F1 | 1641.770 |  | Galactosylated, Core fucosylated |
|  | NSC | H5N2 | 1454.685 |  | Oligomannose |
| 15 | iPSC | H5N2Phos | 1534.675 |  | Phosphorylated, Oligomannose |
|  | NSC | H5N2F1 | 1600.743 |  | Core fucosylated, Oligomannose |
| 16 | H6N2 |  | 1616.738 |  | Oligomannose |
| 17 | D21 iPSC | H6N2 | 1616.738 |  | Oligomannose |
|  | T21 iPSC | H4N3Sia1 | 1786.812 |  | Sialylated |
|  | NSC | H5N4F1 | 2006.902 |  | Core fucosylated,<br>Galactosylated |
| 18 | iPSC | H4N3F1Sia1 | 1932.870 |  | Core fucosylated,<br>Sialylated |
|  | NSC | H5N3F1 | 1803.823 |  | Antennary<br>fucosylated,<br>Agalactosylated,<br>Hybrid |
| 19 | iPSC | H5N5F1 | 2209.981 |  | Core fucosylated,<br>Galactosylated |
|  | NSC | H4N5F2 | 2193.986 |  | Core fucosylated,<br>Antennary<br>fucosylated,<br>Galactosylated |
| 20 | H7N2 |  | 1778.791 |  | Oligomannose |
| 21 | iPSC | H7N2 | 1778.791 |  | Oligomannose |
|  | NSC | H5N3F1Sia1 | 2094.923 |  | Sialylated,<br>Core fucosylated |
| 22 | H6N2Phos |  | 1696.718 |  | Phosphorylated,<br>Oligomannose |
| 23 | H6N3F1 |  | 1965.875 |  | Antennary<br>fucosylated,<br>Galactosylated,<br>Hybrid |
| 24 | H5N4Sia1 |  | 2151.944 |  | Sialylated |
| 25 | iPSC | H8N2 | 1940.844 |  | Oligomannose |
|  | NSC | H5N5F2 | 2356.039 |  | Core fucosylated,<br>Antennary<br>fucosylated,<br>Galactosylated |
| 26 | H8N2 |  | 1940.844 |  | Oligomannose |
| 27 | H5N4F1Sia1 |  | 2298.002 |  | Sialylated,<br>Core fucosylated |
| 28 | H9N2 |  | 2102.897 |  | Oligomannose |
| 29 | H5N4Sia2 |  | 2443.044 |  | Sialylated |
| 30 | H10N2 |  | 2264.949 |  | Oligomannose,<br>Glucosylated |
| 31 | iPSC | H12N2 | 2589.055 |  | Oligomannose,<br>Glucosylated |
|  | NSC | X | X | X | X |

### N-glycosylation of iPSCs vs NSCs

Analysis of the iPSC N-glycome revealed a glycoprofile dominated by four large peaks all belonging to the oligomannose series of N-glycan structures, ranging from M6 to M9 (**Figure 1A, Figure 2**). Consequently, the bulk of the iPSC N-glycome is made up of mannosidic N-glycans (77.92%), with a smaller proportion of complex (16.82%) and hybrid N-glycans (1.57%). A portion of the iPSC N-glycome (3.69%) was taken up by glycans that could not be sorted into any of these basic N-glycan groups and were thus grouped into their own separate group called pseudohybrid glycans (also called monoantennary complex glycans and truncated glycans in literature) (**Figure 2**). Overall, these data indicate that iPSCs possess an extraordinarily basic N-glycosylation pattern which is consistent with the findings of others (Hasehira et al., 2012; Heiskanen et al., 2009; Hemmoranta et al., 2007; Kimura et al., 2021).

**Figure 2.**
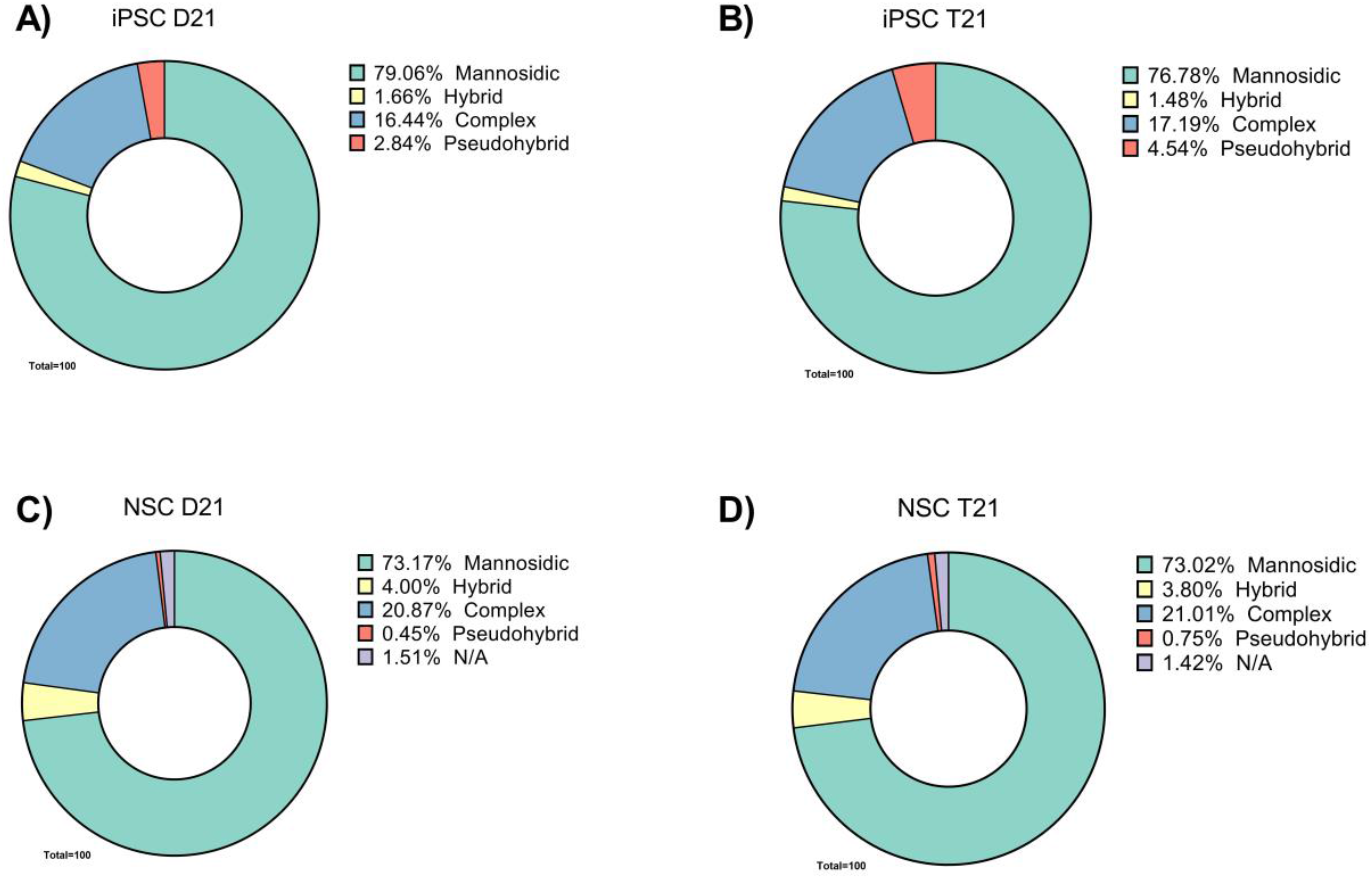
Composition of N-glycomes of induced pluripotent stem cells (iPSCs) and neural stem cells (NSCs) with two (D21)/three (T21) copies of chromosome 21 by N-glycan class. A) iPSC with D21, B) iPSC with T21, C) NSC with D21, D) NSC with T21. Each of the classes is specified in the legend, and N/A refers to GP31 in NSCs which did not have a detectable glycan signal and can therefore not be attributed to any N-glycan class.

Similarly to the iPSCs, NSCs also exhibited a relatively immature N-glycophenotype (**Figure 1B, Figure 2**), with a majority of their N-glycans being mannosidic as well (73.09%), while a smaller part consisted of complex (20.94%) and hybrid (3.90%) N-glycans. Nevertheless, we noted several significant qualitative and quantitative differences between the two stem cell types.

### N-glycosylation of iPSCs vs NSCs – qualitative differences

Firstly, though the chromatograms obtained in the UHPLC analysis looked comparable between iPSCs and NSCs, after MS/MS analysis some differences in the composition of the integrated glycan peaks (GPs) were detected (**Figure 1A, Figure 1B, Table 1**). The most abundant structures were different between iPSCs and NSCs in nine out of 31 total integrated GPs (GP9, GP14, GP15, GP17, GP18, GP19, GP21, GP25, GP31; **Table 1**). The general tendency was for simpler, oligomannose-type glycans to be more present in iPSCs, and other, more complex types of glycans in NSCs. For example, GP21 and GP25 are oligomannose glycans in iPSCs, while in NSCs these peaks contain fucosylated (GP25) and sialylated (GP21) complex type glycans (**Table 1, Figure 1A, Figure 1B, Supplementary Figure 2**).

### N-glycosylation of iPSCs vs NSCs – quantitative differences

The abundances of 17 individual GPs were different between iPSCs and NSCs, 15 of which remained significant after adjusting for multiple testing. Out of those 15 GPs, eight were significantly higher (GP2, GP13, GP15, GP16, GP17, GP19, GP22, GP25), and seven significantly lower (GP12, GP20, GP21, GP26, GP27, GP30, GP31) in NSCs compared to iPSCs (**Table 2, Supplementary Figure 2**). Interestingly, a decrease in Man7 (GP20) and Man8 (GP26) structures previously observed during human iPSC differentiation (Konze et al., 2017) was replicated here (**Table 2, Supplementary Figure 2**).

**Table 2.**
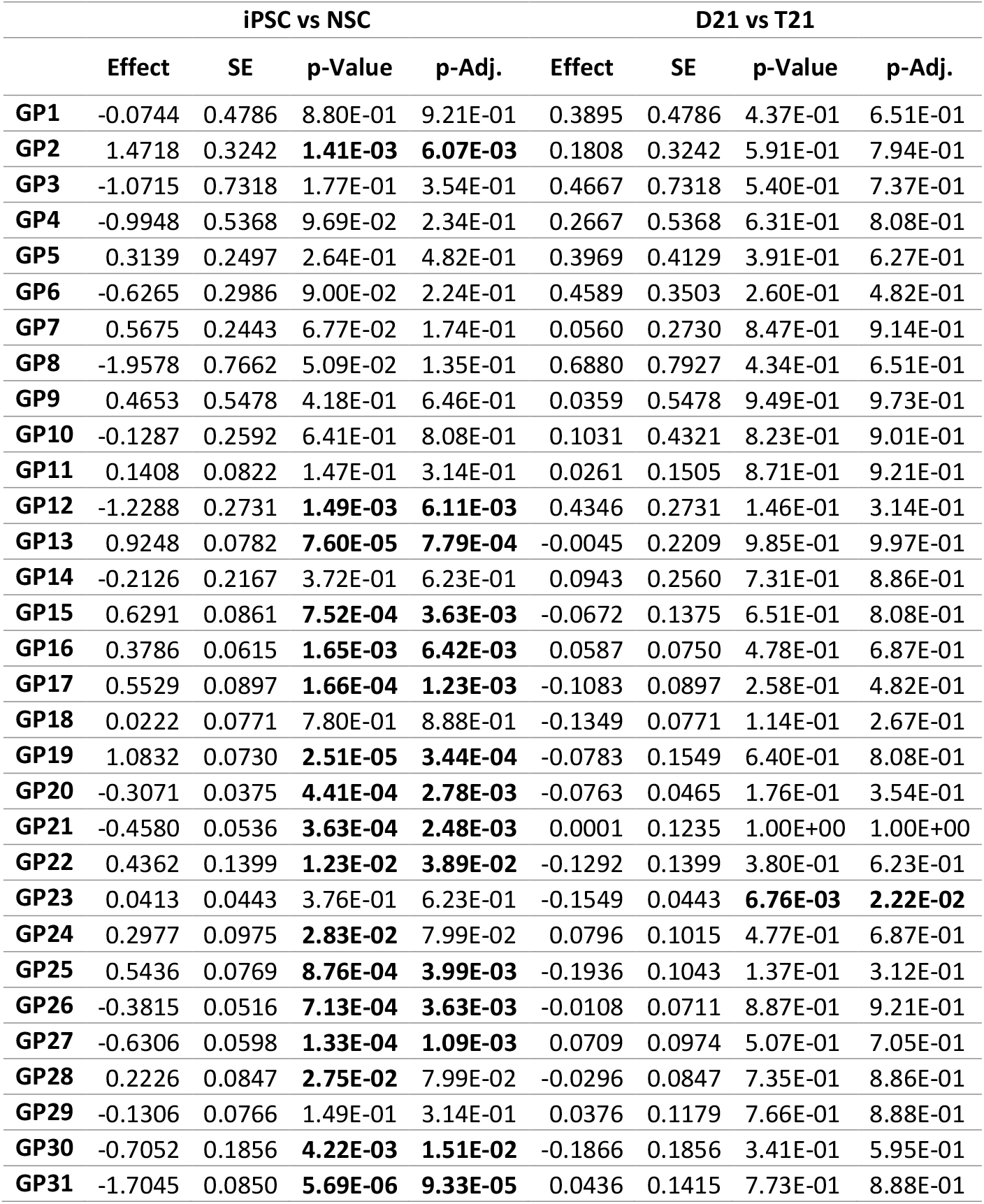
Statistical analysis of individual chromatographic peaks in D21 and T21 iPSCs and NSCs. Significant p-values (<0.05) are indicated in bold font. In iPSCs vs NSCs, the effect value is negative when the chromatographic peak is smaller in NSCs compared to iPSCs, and in D21 vs T21 the effect value is negative when the chromatographic peak is smaller in T21 compared to D21. Association analyses between trisomy status (D21/T21) and differentiation status (iPSC/NSC) and glycomic data were performed using a linear mixed model, and the false discovery rate was controlled using the Benjamini–Hochberg procedure. GP–glycan peak, SE–standard error, iPSC–induced pluripotent stem cells, NSC– neural stem cells, D21–disomy 21, T21–trisomy 21.

|  | iPSC vs NSC |  |  |  | D21 vs T21 |  |  |  |
| --- | --- | --- | --- | --- | --- | --- | --- | --- |
|  | Effect | SE | p-Value | p-Adj. | Effect | SE | p-Value | p-Adj. |
| GP1 | -0.0744 | 0.4786 | 8.80E-01 | 9.21E-01 | 0.3895 | 0.4786 | 4.37E-01 | 6.51E-01 |
| GP2 | 1.4718 | 0.3242 | **1.41E-03** | **6.07E-03** | 0.1808 | 0.3242 | 5.91E-01 | 7.94E-01 |
| GP3 | -1.0715 | 0.7318 | 1.77E-01 | 3.54E-01 | 0.4667 | 0.7318 | 5.40E-01 | 7.37E-01 |
| GP4 | -0.9948 | 0.5368 | 9.69E-02 | 2.34E-01 | 0.2667 | 0.5368 | 6.31E-01 | 8.08E-01 |
| GP5 | 0.3139 | 0.2497 | 2.64E-01 | 4.82E-01 | 0.3969 | 0.4129 | 3.91E-01 | 6.27E-01 |
| GP6 | -0.6265 | 0.2986 | 9.00E-02 | 2.24E-01 | 0.4589 | 0.3503 | 2.60E-01 | 4.82E-01 |
| GP7 | 0.5675 | 0.2443 | 6.77E-02 | 1.74E-01 | 0.0560 | 0.2730 | 8.47E-01 | 9.14E-01 |
| GP8 | -1.9578 | 0.7662 | 5.09E-02 | 1.35E-01 | 0.6880 | 0.7927 | 4.34E-01 | 6.51E-01 |
| GP9 | 0.4653 | 0.5478 | 4.18E-01 | 6.46E-01 | 0.0359 | 0.5478 | 9.49E-01 | 9.73E-01 |
| GP10 | -0.1287 | 0.2592 | 6.41E-01 | 8.08E-01 | 0.1031 | 0.4321 | 8.23E-01 | 9.01E-01 |
| GP11 | 0.1408 | 0.0822 | 1.47E-01 | 3.14E-01 | 0.0261 | 0.1505 | 8.71E-01 | 9.21E-01 |
| GP12 | -1.2288 | 0.2731 | **1.49E-03** | **6.11E-03** | 0.4346 | 0.2731 | 1.46E-01 | 3.14E-01 |
| GP13 | 0.9248 | 0.0782 | **7.60E-05** | **7.79E-04** | -0.0045 | 0.2209 | 9.85E-01 | 9.97E-01 |
| GP14 | -0.2126 | 0.2167 | 3.72E-01 | 6.23E-01 | 0.0943 | 0.2560 | 7.31E-01 | 8.86E-01 |
| GP15 | 0.6291 | 0.0861 | **7.52E-04** | **3.63E-03** | -0.0672 | 0.1375 | 6.51E-01 | 8.08E-01 |
| GP16 | 0.3786 | 0.0615 | **1.65E-03** | **6.42E-03** | 0.0587 | 0.0750 | 4.78E-01 | 6.87E-01 |
| GP17 | 0.5529 | 0.0897 | **1.66E-04** | **1.23E-03** | -0.1083 | 0.0897 | 2.58E-01 | 4.82E-01 |
| GP18 | 0.0222 | 0.0771 | 7.80E-01 | 8.88E-01 | -0.1349 | 0.0771 | 1.14E-01 | 2.67E-01 |
| GP19 | 1.0832 | 0.0730 | **2.51E-05** | **3.44E-04** | -0.0783 | 0.1549 | 6.40E-01 | 8.08E-01 |
| GP20 | -0.3071 | 0.0375 | **4.41E-04** | **2.78E-03** | -0.0763 | 0.0465 | 1.76E-01 | 3.54E-01 |
| GP21 | -0.4580 | 0.0536 | **3.63E-04** | **2.48E-03** | 0.0001 | 0.1235 | 1.00E+00 | 1.00E+00 |
| GP22 | 0.4362 | 0.1399 | **1.23E-02** | **3.89E-02** | -0.1292 | 0.1399 | 3.80E-01 | 6.23E-01 |
| GP23 | 0.0413 | 0.0443 | 3.76E-01 | 6.23E-01 | -0.1549 | 0.0443 | **6.76E-03** | **2.22E-02** |
| GP24 | 0.2977 | 0.0975 | **2.83E-02** | 7.99E-02 | 0.0796 | 0.1015 | 4.77E-01 | 6.87E-01 |
| GP25 | 0.5436 | 0.0769 | **8.76E-04** | **3.99E-03** | -0.1936 | 0.1043 | 1.37E-01 | 3.12E-01 |
| GP26 | -0.3815 | 0.0516 | **7.13E-04** | **3.63E-03** | -0.0108 | 0.0711 | 8.87E-01 | 9.21E-01 |
| GP27 | -0.6306 | 0.0598 | **1.33E-04** | **1.09E-03** | 0.0709 | 0.0974 | 5.07E-01 | 7.05E-01 |
| GP28 | 0.2226 | 0.0847 | **2.75E-02** | 7.99E-02 | -0.0296 | 0.0847 | 7.35E-01 | 8.86E-01 |
| GP29 | -0.1306 | 0.0766 | 1.49E-01 | 3.14E-01 | 0.0376 | 0.1179 | 7.66E-01 | 8.88E-01 |
| GP30 | -0.7052 | 0.1856 | **4.22E-03** | **1.51E-02** | -0.1866 | 0.1856 | 3.41E-01 | 5.95E-01 |
| GP31 | -1.7045 | 0.0850 | **5.69E-06** | **9.33E-05** | 0.0436 | 0.1415 | 7.73E-01 | 8.88E-01 |

Of the ten derived glycan traits that were calculated and compared, five were significantly higher (H, C, GlcNAc, G, F), while the remaining five were significantly lower (M, PH, Phos, Glc, S) in NSCs than iPSCs (**Table 3, Figure 3**). As with the qualitative differences, the general tendency here was for a higher relative amount of hybrid and complex glycans (H, C) and a lower amount of mannosidic glycans (M, Phos, Glc) in NSCs compared to iPSCs.

**Table 3.** Statistical analysis of derived glycan traits in D21 and T21 iPSCs and NSCs. Significant p-values (<0.05) are indicated in bold font. In iPSCs vs NSCs, the effect value is negative when the chromatographic peak is smaller in NSCs compared to iPSCs, and in D21 vs T21 the effect value is negative when the chromatographic peak is smaller in T21 compared to D21. Association analyses between trisomy status (D21/T21) and differentiation status (iPSC/NSC) and glycomic data were performed using a linear mixed model, and the false discovery rate was controlled using the Benjamini–Hochberg procedure. M—mannosidic (pauci- and oligomannose) glycans, H—hybrid glycans, C— complex glycans, PH—pseudohybrid glycans, Phos—phosphorylated glycans, Glc—glycans containing glucose, GlcNAc—glycans with a terminal N-acetylglucosamine, G—glycans containing one or more terminal galactoses, F—fucosylated glycans, S—sialylated glycans, SE–standard error, iPSC–induced pluripotent stem cells, NSC–neural stem cells, D21–disomy 21, T21–trisomy 21.

| Derived trait | iPSC vs NSC |  |  |  | D21 vs T21 |  |  |  |
| --- | --- | --- | --- | --- | --- | --- | --- | --- |
|  | Effect | SE | p-Value | p-Adj. | Effect | SE | p-Value | p-Adj. |
| **M** | -0.0920 | 0.0122 | **6.43E-04** | **3.63E-03** | -0.0227 | 0.0240 | 3.98E-01 | 6.27E-01 |
| **H** | 1.3126 | 0.0481 | **5.75E-10** | **4.72E-08** | -0.1200 | 0.0481 | **3.41E-02** | 9.33E-02 |
| **C** | 0.3175 | 0.0300 | **1.31E-04** | **1.09E-03** | 0.0359 | 0.0717 | 6.42E-01 | 8.08E-01 |
| **PH** | -2.6527 | 0.1841 | **1.60E-07** | **6.57E-06** | 0.6810 | 0.1841 | **4.93E-03** | **1.69E-02** |
| **Phos** | -0.9904 | 0.1970 | **7.11E-04** | **3.63E-03** | 0.0603 | 0.1970 | 7.66E-01 | 8.88E-01 |
| **Glc** | -2.1486 | 0.1673 | **4.31E-07** | **1.18E-05** | -0.1179 | 0.1673 | 4.99E-01 | 7.05E-01 |
| **GlcNAc** | 0.9762 | 0.2688 | **1.50E-02** | **4.56E-02** | 0.0705 | 0.2927 | 8.21E-01 | 9.01E-01 |
| **G** | 1.5882 | 0.0647 | **2.09E-06** | **4.28E-05** | -0.1007 | 0.0743 | 2.47E-01 | 4.82E-01 |
| **F** | 0.7602 | 0.0594 | **5.18E-05** | **6.07E-04** | -0.0183 | 0.0774 | 8.24E-01 | 9.01E-01 |
| **S** | -0.3961 | 0.0657 | **1.80E-03** | **6.71E-03** | 0.1202 | 0.0950 | 2.75E-01 | 4.89E-01 |

**Figure 3.**
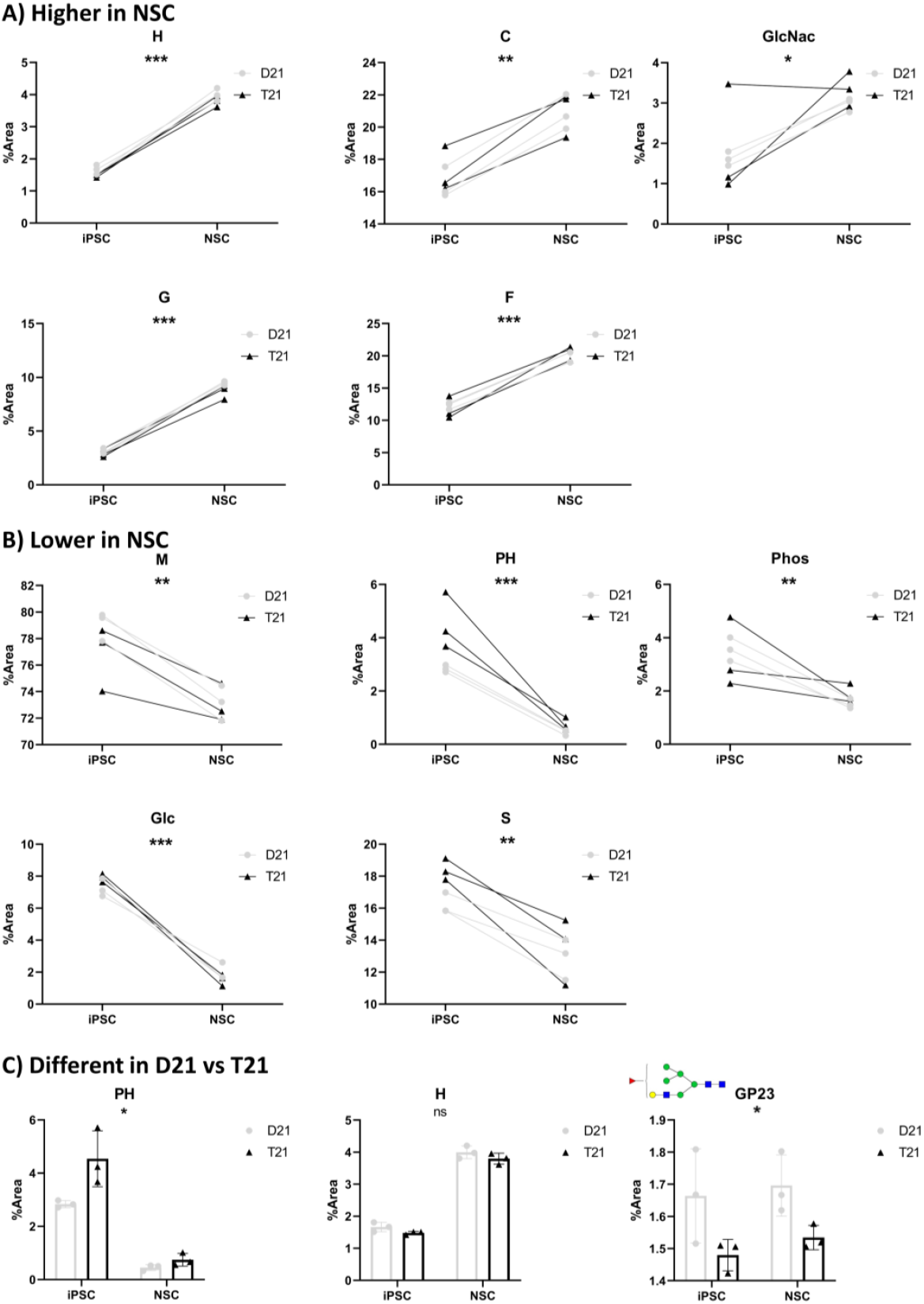
Glycan traits that were shown to be different between either differentiation status or trisomy 21 status. A) Derived traits which are more abundant in NSCs than in iPSCs, B) Derived traits which are less abundant in NSCs than iPSCs, C) Derived traits and one individual glycan peak that are different in D21 vs T21. Each panel shows the %Area value of a different derived glycan trait for each of the analyzed samples. M—mannosidic glycans, H—hybrid glycans, C—complex glycans, PH—pseudohybrid glycans, Phos—phosphorylated glycans, Glc—glycans containing glucose, GlcNAc—glycans with a terminal N-acetylglucosamine, G—glycans containing one or more terminal galactoses, F—fucosylated glycans, S—sialylated glycans. iPSCs are shown first on the x-axis, followed by NSCs. Each clone is represented by two points connected by a line. The clones that have two copies of chromosome 21 (D21) are shown as light gray dots, and those with trisomy 21 (T21) as black triangles. The significance level is marked as follows: ns for nominal significance, one asterisk (*) for p-value < 0.05, two asterisks (**) for p-value < 0.01, and three asterisks (***) for p-value < 0.001.

Overall, our data indicates that the transition from iPSCs to NSCs is accompanied by major changes in the global N-glycosylation profile characterized by a shift from simple, immature N-glycans towards more complex and elaborate ones.

### N-glycosylation in an isogenic model of trisomy 21 (D21 vs T21)

When comparing D21 and T21 clones, the differences were expectedly more subtle than between cell types. Still, some notable qualitative and quantitative differences indicating the effect of T21 on total cell N-glycosylation were observed.

### N-glycosylation in an isogenic model of trisomy 21 (D21 vs T21) – qualitative differences

A difference in composition between D21 and T21 was observed only in GP17, which is an oligomannose-type glycan (H6N2) in D21 iPSCs, and a monoantennary sialylated complex glycan (H4N3Sia1) in T21 iPSCs. The composition of GP17 also differs between iPSCs and NSCs (**Table 1, Figure 1C**).

### N-glycosylation in an isogenic model of trisomy 21 (D21 vs T21) – quantitative differences

The only individual glycan peak that was different between D21 and T21 was GP23, a fucosylated hybrid glycan (H6N3F1) significantly less abundant in T21 cells (**Table 2, Figure 3C**). Regarding derived glycan traits, pseudohybrid glycans (PH) were significantly higher, and hybrid glycans (H) nominally lower in T21 cells (**Table 3**). While the nominal decrease in H was most likely driven by the change in GP23 (seeing as GP23 is a hybrid glycan), the significant increase in PH, more significantly pronounced in iPSCs than NSCs (**Figure 2**) is an interesting observation without clear functional implications as of yet.

To sum up, the differences in N-glycosylation between D21 and T21 isogenic clones are less pronounced than those observed when comparing iPSCs and NSCs. Still, they indicate that the presence of an extra copy of chromosome 21 clearly causes a shift in N-glycosylation that can be traced back to the earliest developmental stages.

## Discussion

We have conducted an analysis of the total N-glycome in an isogenic cell line model of trisomy 21 (T21) at two different stages of differentiation, iPSCs and NSCs. These cell lines are a convenient *in vitro* model for T21 research because they were obtained from a single person with mosaic Down syndrome (DS), meaning they are otherwise genetically identical (Murray et al., 2015).

### Cell type differences

We first compared total cell N-glycosylation between iPSCs and NSCs, where notable differences in the N-glycomes of different cell types were observed. This is in line with a recent study that showed significant differences in glycopeptide composition using MS analysis during human neuronal differentiation starting from iPSCs (Cindrić et al., 2021; Kimura et al., 2021; Nižetić & Groet, 2012). A glycan structure that they describe and mainly focus on, previously described in the context of cell glycans as a brain-only glycan, H5N3F1, has not been detected in our cells. This is in agreement with their research as this structure was only detected in cells that have differentiated past the NSC stage into neuron-like cells, so we would not expect to detect it in our NSCs (Kimura et al., 2021; Shimizu et al., 1993). Alternatively (but less likely) some differences between the two systems might arise from the different method used to reprogramme the primary cells into iPSCs: while Kimura et al. used an iPSC line generated using lentiviral (integrational) approach (Kimura et al., 2021), our iPSC lines were generated using Sendai virus (non-integrational) re-programming (Murray et al., 2015).

We show differences in abundance in a large fraction of the individual glycan peaks and all of the studied derived glycan traits between iPSCs and NSCs, showing a shift in the cell N-glycosylation profile. These findings add further knowledge to the previously observed differential glycosylation during the differentiation of other cell types and tissues (Link-Lenczowski et al., 2019; Liu et al., 2018; Yale et al., 2018).

As theoretically expected, our data find the levels of more immature, evolutionarily older glycans (i.e. mannosidic glycans) higher in iPSCs, whereas the levels of hybrid and complex glycans are generally higher in NSCs. This finding replicates previous research wherein human iPSCs show a significantly higher content of mannosidic glycans when compared to more differentiated cells (Hasehira et al., 2012; Heiskanen et al., 2009; Hemmoranta et al., 2007) and can be explained by the higher level of stemness of iPSCs, which are less specialized than NSCs and thus express a different protein repertoire, with less specialized, neuron-specific proteins with niche roles that require for them to be specifically glycosylated (Scott & Panin, 2014; Varderidou-Minasian et al., 2020). All of these findings confirm the validity of our cell-line models and used methods.

Interestingly, an oligomannose structure containing three glucose residues (Glc3-Man9-GlcNAc2) was only observed in iPSCs (GP31) and was either not present at all or present in quantities below the level of detection in NSCs. This structure is the precursor of all N-glycans and is transferred from a dolichol phosphate anchor to a nascent polypeptide in the endoplasmic reticulum. After the glycan has been transferred to the protein, all of the glucose residues and one of the mannose residues get enzymatically cleaved, after which the protein is further transferred for sequential processing. This process also serves as a quality control checkpoint of protein folding, meaning that all of the polypeptides containing this glycan structure are yet to be submitted to quality control and are still considered immature (Parodi, 2000; Stanley et al., 2022). Given that it was only detected in iPSCs, the presence of this glycan is potentially another marker of cell stemness.

### D21 vs T21

In our comparison of isogenic cells with and without trisomy 21, we uncover subtle, yet notable differences. In one specific individual glycan peak, there is a difference in proposed composition already at the iPSC stage. Namely, GP17 contains an oligomannose-type glycan (H6N2) in D21 iPSCs, while in T21 iPSCs the most abundant structure of GP17 is a monoantennary sialylated glycan (H4N3Sia1). While not a major change, this complex type of glycan being more dominant in one GP in T21 agrees with the phenomenon of premature aging in DS (Cindrić et al., 2021; Franceschi et al., 2019; Nižetić & Groet, 2012) and suggests that it might start on a subcellular level even earlier than previously thought.

One GP whose abundance is significantly higher in D21 than T21 and remains so at both examined stages of differentiation is H6N3F1 (GP23), a monoantennary hybrid glycan with an antennary fucose. This glycan has previously been reported in multiple human tissues— blood plasma, more specifically human coagulation factor V (Ma et al., 2020), prefrontal cortex (Lee et al., 2020) and adipose-derived stem cells (Wongtrakul-Kish et al., 2021). However, to the best of our knowledge, it has not been reported to change in a significant manner in any physiological condition or disease. This finding potentially makes this glycan interesting, as it seems to be uniquely decreased in T21. Moreover, it would be interesting to further explore whether this difference functionally affects the NSCs, or their cell fate in further differentiation, as N-glycosylation differences have been observed associated with such changes (Yale et al., 2018), and quantitative cell fate differences during neural differentiation are associated with DS (Huo et al., 2018; Olmos-Serrano et al., 2016).

We also observed a nominally significant increase in the relative amount of hybrid glycans in D21 as opposed to T21, driven by the aforementioned significant increase in H6N3F1 (GP23), which is a hybrid glycan. Additionally, pseudohybrid glycans (sometimes also referred to as monoantennary complex glycans and truncated glycans) were significantly higher in T21 cells, especially so in T21 iPSCs. In the adult brain, there are putative pathways involved in the biosynthesis of these glycans mostly based on the activity of N-acetyl-D-hexosaminidase B, an enzyme encoded by *HEXB* (Klarić & Lauc, 2022). However, this gene is located on chromosome 5, so the implications or causes of this observation are yet to be elucidated in the case of our study.

Our statistical model considered all samples at once to gain more statistical power. However, as the sample number was low, we also looked at the values individually for each clone in both sample types and saw a trend in some of the analyzed traits. Perhaps most interestingly, we noticed all of the D21 iPSC clones had lower sialylation than their T21 counterparts (**Figure 3**). While intriguing, the functional meaning of this finding is hard to interpret using our analytical approach, as sialylation changes in either direction can affect cell fate decision (Li & Ding, 2019), depending on the protein carrying the sialylated glycan. It is possible that higher levels of sialic acids in T21 iPSCs could preferentially send them down different differentiation routes, though that hypothesis is yet to be tested.

Inevitably, the small number of samples is a limitation of this study due to the low power of the statistical tests to detect small effects. The analysis was also performed on cells that were all derived from the same person, which, while giving us an ideal platform to study the isolated effect of T21, still limits us in the sense of generalization of our findings. It is also important to note that the changes we detected in N-glycans are potentially affected by modified levels of the corresponding peptide, however we can only speculate on this as our method does not produce glycoproteomic data. A larger study with cell lines derived from multiple people and including multiple different types of analysis (glycoproteomic in addition to UHPLC) would be required to address these issues.

## Methods

### Cell culture

We cultured a total of six iPSC lines, all derived from an adult diagnosed with constitutional mosaicism for DS. Three of the cell lines were disomic for chromosome 21 (D21), and the other three were trisomic for chromosome 21 (T21) while otherwise being genetically identical. The D21 lines are NIZEDSM1iD21-C3, -C7, and -C9 while the T21 lines are NIZEDSM1iT21-C5, -C6, -C13 (Murray et al., 2015). iPSCs were cultured on plates coated with Geltrex in E8 media at 37°C, 5% CO_2_.

From these iPSC lines we generated NSC lines following Gibco Life Techologies protocol (MAN0008031). NSCs were cultured on plates coated with Geltrex in Neural Expansion media at 37°C, 5% CO_2_.

### Cell validation

The cells were validated with following methods: FISH to check trisomy for chromosome 21 in T21 cells and immunostaining. iPSCs were stained for pluripotency markers (OCT4, SSEA4, TRA1-60 and TRA1-81) while the NSCs were stained NSCs specific markers (SOX2, PAX6 and Nestin). Detailed protocol for FISH as well as immunostaining were performed as previously described (Alic et al., 2021).

### N-glycosylation analysis of harvested cells

#### Deglycosylation, fluorescent labelling and clean-up of cell N-glycans

The cells were collected using accutase and pelleted by centrifugation for 5 min at 300 x g. Cell pellets containing around 2×10^6^ cells for iPSCs and around 5×10^6^ cells for NSCs were snap frozen in dry ice and stored at -80°C until ready for analysis.

The cell pellets were thawed before proceeding with a previously described lysate in-solution deglycosylation protocol (Vacchini et al., 2020). Briefly, the cell pellets were lysed and glycoproteins extracted using chloroform and methanol. Enzymatic deglycosylation was done with PNGase F, after which the released N-glycans were fluorescently labelled using procainamide. Excess dye and other reagents were removed using solid phase extraction (SPE) with an Acroprep 1 mL 0.2 μm wwPTFE plate (Pall, NY, USA) used as stationary phase (Pucic et al., 2011; Trbojević Akmačić et al., 2015). Glycan enrichment was done using cotton hydrophilic interaction liquid chromatography (HILIC) SPE (Selman et al., 2011).

#### HILIC-UHPLC-FLR analysis of procainamide-labelled cell N-glycans

Fluorescently labelled cell N-glycans were analyzed by ultra-high performance liquid chromatography based on hydrophilic interactions with fluorescence detection (HILIC-UHPLC-FLR) using an Acquity UPLC H-class instrument controlled by the Empower 3 software, build 3471 (Waters, MA, USA). Before injection the samples were prepared by mixing the purified N-glycan sample (in water) and acetonitrile 27%/73% (v/v) and were kept at 10 °C. Separation of N-glycans was done at 25 °C on an Acquity UPLC Glycan bridged ethylene hybrid (BEH) amide column, 130 Å, 1.7 μm BEH particles, 2.1 mm × 150 mm; Waters, MA, USA). Solvent A was 100 mM ammonium formate, pH 4.4, and solvent B was acetonitrile. The chromatographic run was performed over 95 min at a flow rate of 0.561 mL/min with a linear gradient of 27–38.7% solvent A. The excitation wavelength was set at 310 nm, while the emission wavelength was set at 370 nm. The acquired chromatograms were manually integrated, wherein each chromatogram was separated into 31 N-glycan peaks (GPs; **Figure 1**), producing %Area values for each GP. %Area is the percentage of the total integrated chromatogram area that a certain GP takes up and is representative of the percentage that its corresponding glycan takes up in the analyzed glycome.

#### LC-MS-MS analysis of procainamide-labelled cell N-glycans

N-glycans from each experimental group (i.e. D21 iPSCs, T21 iPSCs, D21 NSCs, and T21 NSCs) were pooled to create a representative sample that was used for qualitative structural analysis by UHPLC coupled to mass spectrometer (LC-MS-MS).

The separation (UHPLC) method was the same as described above, and the mass spectrometer was set up as follows: positive ion mode, ionBooster source, capillary voltage 3.6 kV, dry gas temperature 180°C and flow 4 L/min. The mass-to-charge ratio (m/z) range over which mass spectra were recorded was from 100 to 4000 m/z and spectra were acquired at a frequency of 0.5 Hz. Tandem MS (MS/MS) experiments were performed in the data-dependent acquisition mode with the three precursors of highest intensity being selected to undergo CID (collision-induced dissociation) fragmentation. The m/z range for MS/MS spectra was 100–4000 m/z. Spectra were processed in Bruker Compass Data Analysis 4.4.

The acquired data was analysed manually, wherein glycan compositions were proposed via searching GlycoMod (http://web.expasy.org/glycomod, Cooper et al., 2001) for monoisotopic masses of the ions detected in the base peak chromatogram, with mass tolerance of ±0.4 Daltons. Glycan compositions and structures were manually confirmed using retention times and tandem MS fragmentation data which was searched for diagnostic product ions. The proposed structures were then depicted using GlycoWorkbench 2.1 stable build 146 (Ceroni et al., 2008).

Derived traits, used to group glycans with the same compositional/structural features, were calculated from the %Area values of directly measured GPs. A total of ten derived traits were calculated, namely M—mannosidic (pauci- and oligomannose) glycans, H—hybrid glycans, C—complex glycans, PH—pseudohybrid glycans, Phos—phosphorylated glycans, Glc—glycans containing glucose, GlcNAc—glycans with a terminal N-acetylglucosamine, G— glycans containing one or more terminal galactoses, F—fucosylated glycans, S—sialylated glycans (**Supplementary Table 2**).

### Statistical analysis

Association analyses between trisomy status (D21/T21) and differentiation status (iPSC/NSC) and glycomic data were performed using a linear mixed model. Analyses included glycan measurement as a dependent continuous variable, individual identification (ID) was included in a model as a random intercept, and trisomy 21 and cell differentiation status were included as fixed effects. False discovery rate was controlled using the Benjamini–Hochberg procedure (function p.adjust(method = “BH”)). Data were analyzed using R programming language (version 4.0.2).

