## Supplemental information for "Total cell N-glycosylation is altered during neuronal differentiation of iPSC to NSC and is disturbed by trisomy 21"

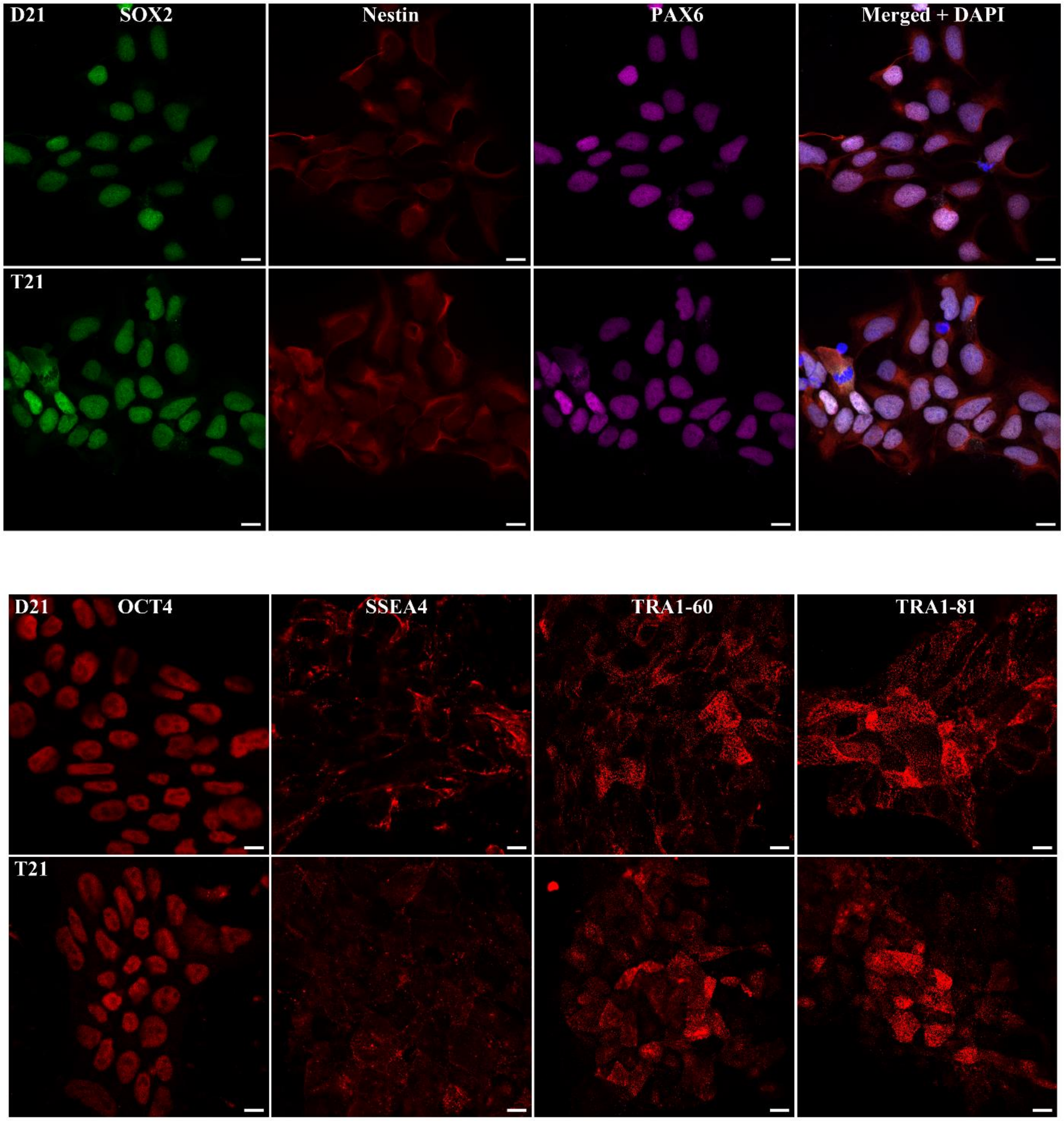


**Supplementary Figure 1. Cellular validation.** A) iPSCs were validated by immunostaining to confirm the expression of a panel of pluripotency markers: OCT4, SSEA4, TRA1-60 and TRA1-81. Scale bar 10 µm. B) NSCs were validated by immunostaining to confirm the expression of Nestin, SOX2 and PAX6. Scale bar 10 µm.

**Supplementary Table 1. Theoretical and observed masses of most abundant glycans for each peak and each sample.** H – hexose, N – N-acetylglucosamine, F – fucose, Phos – phosphate, Sia – sialic acid, iPSC – induced pluripotent stem cells, NSC – neural stem cells, D21 – disomic cells, T21 – cells with three copies of chromosome 21.

| GP | Composition | Theoretical mass | iPSC D21 Observed mass | | iPSC T21 Observed mass | | NSC D21 Observed mass | | NSC T21 Observed mass | |
|---|---|---|---|---|---|---|---|---|---|---|
| | | [M:ProA+H+]+ | m/z | Delta | m/z | Delta | m/z | Delta | m/z | Delta |
| 1 | H3N2 | 1130.5797 | 1130.468 | 0.1119 | 1130.469 | 0.1107 | 1130.462 | 0.1175 | 1130.463 | 0.1166 |
| 2 | H3N2F1 | 1276.6376 | 1276.519 | 0.1185 | 1276.521 | 0.1170 | 1276.510 | 0.1274 | 1276.515 | 0.1229 |
| 3 | H3N3 | 1333.6591 | 1333.546 | 0.1127 | 1333.547 | 0.1124 | 1333.528 | 0.1308 | 1333.537 | 0.1219 |
| 4 | H3N2P1 | 1210.5691 | 1210.465 | 0.1045 | 1210.462 | 0.1074 | 1210.460 | 0.1094 | 1210.458 | 0.1111 |
| 5 | H4N2 | 1292.6325 | 1292.517 | 0.1160 | 1292.508 | 0.1246 | 1292.508 | 0.1250 | 1292.507 | 0.1252 |
| 6 | H4N2 | 1292.6325 | 1292.515 | 0.1180 | 1292.511 | 0.1217 | 1292.513 | 0.1191 | 1292.506 | 0.1266 |
| 7 | H3N3F1 | 1479.7170 | 1479.589 | 0.1284 | 1479.589 | 0.1279 | 1479.577 | 0.1400 | 1479.583 | 0.1343 |
| 8 | H3N4 | 1536.7385 | 1536.601 | 0.1380 | 1536.597 | 0.1419 | 1536.612 | 0.1268 | 1536.603 | 0.1353 |
| 9 | H4N2F1 | 1438.6905 | | | | | 1438.561 | 0.1298 | 1438.562 | 0.1289 |
| | H5N2P1 | 1534.6747 | 1534.543 | 0.1321 | 1534.532 | 0.1432 | | | | |
| 10 | H4N2P1 | 1372.6219 | 1372.513 | 0.1093 | 1372.507 | 0.1147 | 1372.505 | 0.1170 | 1372.504 | 0.1180 |
| 11 | H3N4F1 | 1682.7964 | 1682.666 | 0.1300 | 1682.656 | 0.1403 | 1682.657 | 0.1391 | 1682.655 | 0.1414 |
| 12 | H5N2 | 1454.6854 | 1454.562 | 0.1231 | 1454.559 | 0.1266 | 1454.551 | 0.1344 | 1454.556 | 0.1291 |
| 13 | H5N2 | 1454.6854 | 1454.561 | 0.1246 | 1454.558 | 0.1246 | 1454.553 | 0.1322 | 1454.552 | 0.1334 |
| 14 | H5N2 | 1454.6854 | | | | | 1454.556 | 0.1290 | 1454.552 | 0.1335 |
| | H4N3F1 | 1641.7698 | 1641.642 | 0.1280 | 1641.634 | 0.1358 | | | | |
| 15 | H5N2P1 | 1534.6747 | 1534.559 | 0.1156 | 1534.549 | 0.1253 | | | | |
| | H5N2F1 | 1600.7433 | | | | | 1600.608 | 0.1349 | 1600.604 | 0.1397 |
| 16 | H6N2 | 1616.7382 | 1616.607 | 0.1311 | 1616.602 | 0.1358 | 1616.601 | 0.1368 | 1616.597 | 0.1411 |
| 17 | H6N2 | 1616.7382 | 1616.606 | 0.1323 | | | | | | |
| | H4N3Sia1 | 1786.8119 | | | 1786.675 | 0.1372 | | | | |
| | H5N4F1 | 2006.9020 | | | | | 2006.759 | 0.1431 | 2006.750 | 0.1524 |
| 18 | H5N3F1 | 1803.8226 | | | | | 1803.681 | 0.1417 | 1803.678 | 0.1443 |
| | H4N3F1Sia1 | 1932.8698 | 1932.729 | 0.1408 | 1932.720 | 0.1497 | | | | |
| 19 | H4N5F2 | 2193.9865 | | | | | 2193.838 | 0.1487 | 2193.827 | 0.1594 |
| | H5N5F1 | 2209.9814 | 2209.838 | 0.1436 | 2209.841 | 0.1401 | | | | |
| 20 | H7N2 | 1778.7910 | 1778.652 | 0.1389 | 1778.649 | 0.1417 | 1778.647 | 0.1445 | 1778.644 | 0.1471 |
| 21 | H7N2 | 1778.7910 | 1778.650 | 0.1412 | 1778.645 | 0.1460 | | | | |
| | H5N3F1Sia1 | 2094.9226 | | | | | 2094.770 | 0.1525 | 2094.768 | 0.1551 |
| 22 | H6N2P1 | 1696.7276 | | | | | 1696.564 | 0.1634 | 1696.557 | 0.1704 |
| 23 | H6N3F1 | 1965.8755 | | | | | 1965.732 | 0.1431 | 1965.729 | 0.1466 |
| 24 | H5N4Sia1 | 2151.9441 | 2151.798 | 0.1457 | 2151.791 | 0.1535 | 2151.791 | 0.1531 | 2151.795 | 0.1490 |
| 25 | H8N2 | 1940.8438 | 1940.705 | 0.1384 | 1940.703 | 0.1408 | | | | |
| | H5N5F2 | 2356.0393 | | | | | 2355.893 | 0.1463 | 2355.885 | 0.1545 |
| 26 | H8N2 | 1940.8438 | 1940.699 | 0.1452 | 1940.694 | 0.1503 | 1940.691 | 0.1531 | 1940.686 | 0.1579 |
| 27 | H5N4F1Sia1 | 2298.0020 | 2297.846 | 0.1557 | 2297.848 | 0.1537 | 2297.830 | 0.1720 | 2297.841 | 0.1613 |
| 28 | H9N2 | 2102.8967 | 2102.745 | 0.1519 | 2102.747 | 0.1501 | 2102.737 | 0.1594 | 2102.733 | 0.1641 |
| 29 | H5N4Sia2 | 2443.0441 | 2442.892 | 0.1523 | 2442.867 | 0.1770 | 2442.887 | 0.1576 | 2442.825 | 0.2193 |
| 30 | H10N2 | 2264.9495 | 2264.806 | 0.1439 | 2264.798 | 0.1520 | 2264.797 | 0.1525 | 2264.801 | 0.1489 |
| 31 | H12N2 | 2589.0551 | 2588.920 | 0.1353 | 2588.928 | 0.1267 | X | X | X | X |

**Supplementary Table 2. Derived traits calculation formulas for each of the analyzed cell types.** M—mannosidic (pauci- and oligomannose) glycans, H—hybrid glycans, C—complex glycans, PH—pseudohybrid glycans, Phos—phosphorylated glycans, Glc—glycans containing glucose, GlcNAc—glycans with a terminal N-acetylglucosamine, G—glycans containing one or more terminal galactoses, F—fucosylated glycans, S—sialylated glycans. There were no differences in the most abundant structures in GPs between D21 and T21 NSCs, so the same formulas were used to calculate their derived traits.

| | |
|---|---|
| **D21 iPSC** | |
| **M** | GP1 + GP2 + GP4 + GP5 + GP6 + GP9 + GP10 + GP12 + GP13 + GP15 + GP16 + GP17 + GP20 + GP21 + GP22 + GP25 + GP26 + GP28 + GP30 + GP31 |
| **H** | GP23 |
| **C** | GP8 + GP11 + GP19 + GP24 + GP27 + GP29 |
| **PH** | GP3 + GP7 + GP14 + GP18 |
| **Glc** | GP30 + GP31 |
| **Phos** | GP4 + GP9 + GP10 + GP15 + GP22 |
| **GlcNAc** | GP3 + GP7 + GP8 + GP11 |
| **G** | GP14 + GP19 + GP23 |
| **F** | GP2 + GP7 + GP11 + GP14 + GP18 + GP19 + GP23 + GP27 |
| **S** | GP18 + GP24 + GP27 + GP29 |
| **T21 iPSC** | |
| **M** | GP1 + GP2 + GP4 + GP5 + GP6 + GP9 + GP10 + GP12 + GP13 + GP15 + GP16 + GP20 + GP21 + GP22 + GP25 + GP26 + GP28 + GP30 + GP31 |
| **H** | GP23 |
| **C** | GP8 + GP11 + GP19 + GP24 + GP27 + GP29 |
| **PH** | GP3 + GP7 + GP14 + GP17 + GP18 |
| **Glc** | GP30 + GP31 |
| **Phos** | GP4 + GP9 + GP10 + GP15 + GP22 |
| **GlcNAc** | GP3 + GP7 + GP8 + GP11 |
| **G** | GP14 + GP19 + GP23 |
| **F** | GP2 + GP7 + GP11 + GP14 + GP18 + GP19 + GP23 + GP27 |
| **S** | GP17 + GP18 + GP24 + GP27 + GP29 |
| **NSC** | |
| **M** | GP1 + GP2 + GP4 + GP5 + GP6 + GP9 + GP10 + GP12 + GP13 + GP14 + GP15 + GP16 + GP20 + GP22 + GP26 + GP28 + GP30 |
| **H** | GP18 + GP21 + GP23 |
| **C** | GP8 + GP11 + GP17 + GP19 + GP24 + GP25 + GP27 + GP29 |
| **PH** | GP3 + GP7 |
| **Glc** | GP30 |
| **Phos** | GP4 + GP10 + GP22 |
| **GlcNAc** | GP3 + GP7 + GP8 + GP11 + GP18 |
| **G** | GP17 + GP19 + GP23 + GP25 |
| **F** | GP2 + GP7 + GP9 + GP11 + GP15 + GP17 + GP18 + GP19 + GP21 + GP23 + GP25 + GP27 |
| **S** | GP21 + GP24 + GP27 + GP29 |

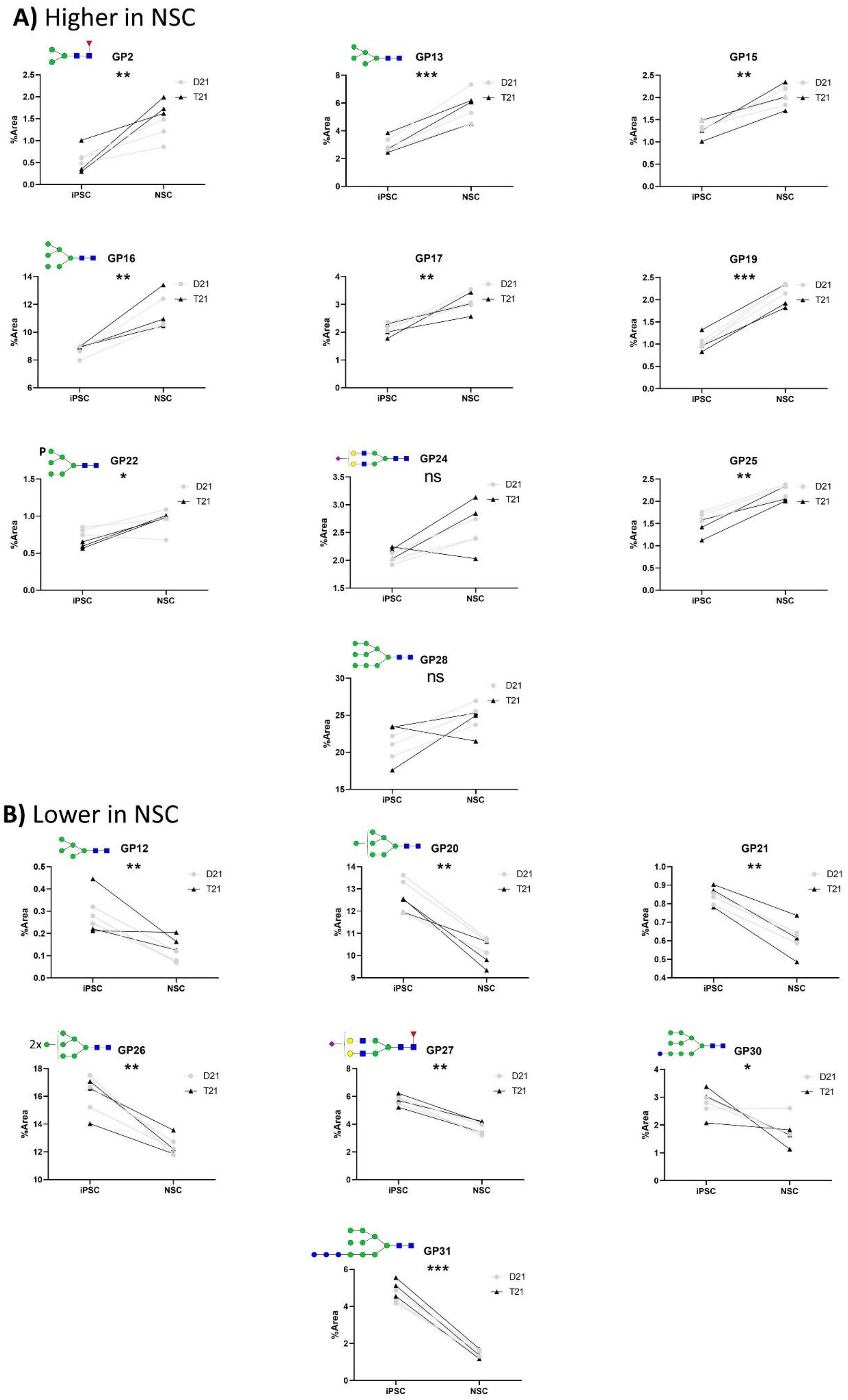
A) Higher in NSC
GP2
**
GP13
***
GP15
**
GP16
**
GP17
**
GP19
***
GP22
*
GP24
ns
GP25
**
GP28
ns
B) Lower in NSC
GP12
**
GP20
**
GP21
**
GP26
**
GP27
**
GP30
*
GP31
***
2x
P
D21
T21
%Area
iPSC
NSC

**Supplementary Figure 2. Individual glycan peaks showing statistically significant differences in abundance between iPSC and NSC.** A) Glycan peaks (GPs) which are more abundant in NSCs than in iPSCs, B) GPs which are less abundant in NSCs than iPSCs. Each panel shows the %Area value of a different GP for each of the analyzed samples. iPSCs are shown first on the x-axis, followed by NSCs. Each clone is represented by two points connected by a line. The clones that have two copies of chromosome 21 (D21) are shown as gray dots, and those with trisomy 21 (T21) as black triangles. For the GPs that have the same most abundant structure in both iPSCs and NSCs, the structure is shown above the corresponding graph. All other peaks are annotated in Main Figure 1. The significance level is marked as follows: ns for nominal significance, one asterisk (*) for p-value < 0.05, two asterisks (**) for p-value < 0.01, and three asterisks (***) for p-value < 0.001.
